# Backbone amides are key determinants of Cl^−^ selectivity in CLC ion channels

**DOI:** 10.1101/2022.07.01.498497

**Authors:** Lilia Leisle, Kin Lam, Sepehr Dehghani-Ghahnaviyeh, Eva Fortea, Jason Galpin, Christopher A. Ahern, Emad Tajkhorshid, Alessio Accardi

## Abstract

Chloride homeostasis is tightly regulated in cellular compartments by dedicated channels and transporters. Whereas CLC-type channels select for Cl^**−**^ over other anions, all other ‘chloride’ channels are indiscriminate in their anionic permeability. Pore-lining side chains are thought to determine Cl^**−**^ selectivity in CLC channels. However, orientation and functional roles of these side chains is not conserved among CLCs. All CLC pores are lined by backbone amides in a conserved structural arrangement, suggesting a role of mainchain groups in selectivity. We replaced pore-lining residues in the CLC-0 and bCLC-k channels with their respective α-hydroxy acid counterparts using nonsense suppression method. This exchanges peptide-bond amides with ester-bond oxygens, incapable of hydrogen-bonding with permeating anions. Backbone substitutions functionally degrade inter-anion discrimination in a site-specific manner. These effects depend on the presence of a glutamate side chain that competes with ions permeating through the pore. Molecular dynamics simulations show that ion energetics within the bCLC-k pore are primarily determined by interactions with backbone amides. Insertion of an α-hydroxy acid significantly alters ion selectivity and global pore hydration. We propose that backbone amides are conserved determinants of Cl^**−**^ specificity in CLC channels in a mechanism reminiscent of that described for K^+^ channels.

## Introduction

Anion-selective channels and transporters control Cl^**−**^ homeostasis in all living cells and within their intracellular compartments. The ability of these channels to select against cations and discriminate amongst physiological anions is central to their function *in vivo*. While cation selectivity mechanisms are relatively well understood ^1, 2, 3, 4, 5, 6, 7, 8^, the principles underlying Cl^**−**^ channel selectivity are poorly resolved. Indeed, most ‘Cl^**−**^ channels’ are more permeable to anions other than their biological namesake: GABA ^9^, CFTR ^10^, TMEM16A ^11^, TMEM16B ^12^, and Bestrophin ^13, 14^ channels follow the Hofmeister lyotropic selectivity sequence ^15^ of SCN^**−**^>I^**−**^>NO_3_^**−**^>Br^**−**^>Cl^**−**^, with slight deviations. In contrast, CLC-type channels and transporters select for Cl^**−**^ over other anions with a sequence of Cl^**−**^>Br^**−**^>NO_3_^**−**^>I^**−**^ ^16, 17, 18, 19, 20, 21^. This selectivity sequence is evolutionarily well-conserved from prokaryotes to eukaryotes and between transporters and channels. Notably, while most CLCs are Cl^**−**^ selective, the atCLC-a exchanger from *Arabidopsis thaliana* is NO_3_^**−**^ selective ^22, 23^, and members of a clade of prokaryotic CLCs are highly F^**−**^ selective ^24, 25, 26, 27^. Thus, the CLC pore provides a unique and plastic structural template to investigate the mechanisms that underlie anion selectivity.

All CLCs are dimers, where each monomer forms a separate Cl^**−**^ permeation pathway ^28, 29^. The CLC-ec1 structure allowed for the identification of three anionic binding sites ^30, 31, 32, 33, 34, 35, 36, 37^, coined S_int_, S_cen_ and S_ext_ for internal, central and external, respectively (Fig. 1A-C), whose position and coordination are evolutionarily conserved in eukaryotic channels and transporters. Coordination with permeant anions at the internal S_int_ site is weak as the dehydration of ions is only partially compensated via interactions with the backbone amides of a serine residue at position 107 (called Ser_cen_) and Glycine 108 (using CLC-ec1 numbering) in the loop connecting helices C and D (Fig. 1A, C) ^19, 30, 38^. Conversely, anions positioned in the central and external sites, S_cen_ and S_ext_, interact with the protein more extensively, consistent with a key role of this region in selectivity. In S_cen_, Cl^−^ is coordinated by the conserved side chains of S107 (Ser_cen_) and of Y445 (Tyr_cen_), as well as the backbone amides of I356 and F357 (Fig. 1A). Ion coordination in S_ext_ is mediated by the backbone amides of E148 (Glu_ex_), G149, F357 and A359 (Fig.1A). Thus, interactions with side chains and backbone amides contribute to the preferential stabilization of anions over cations within the CLC pore ^31, 39^. The pore architecture is conserved in the mammalian bCLC-k and hCLC-1 channels, involving similar coordination patterns (Fig. 1B-C), with the notable exception that in bCLC-k Ser_cen_ points away from S_cen_ (Fig. 1B). The negatively charged side chain of Glu_ex_ is a tethered anion that can occupy the S_cen_ and S_ext_ sites with similar coordination to that of the bound Cl^−^ ions ^31, 32, 39^ (Fig. 1, Supp. Fig. 1). The competition between Glu_ex_ and the Cl^−^ ions is essential for CLC function ^28, 29, 32, 40, 41, 42^ and weakened ion binding at the S_cen_ site alters Cl^−^/H^+^ exchange stoichiometry in the transporters and gating in channels ^17, 20, 43, 44, 45, 46, 47, 48, 49^. Thus, the molecular and energetic determinants of selective anion binding and permeation also govern the CLC transport mechanism.

**Figure 1.**
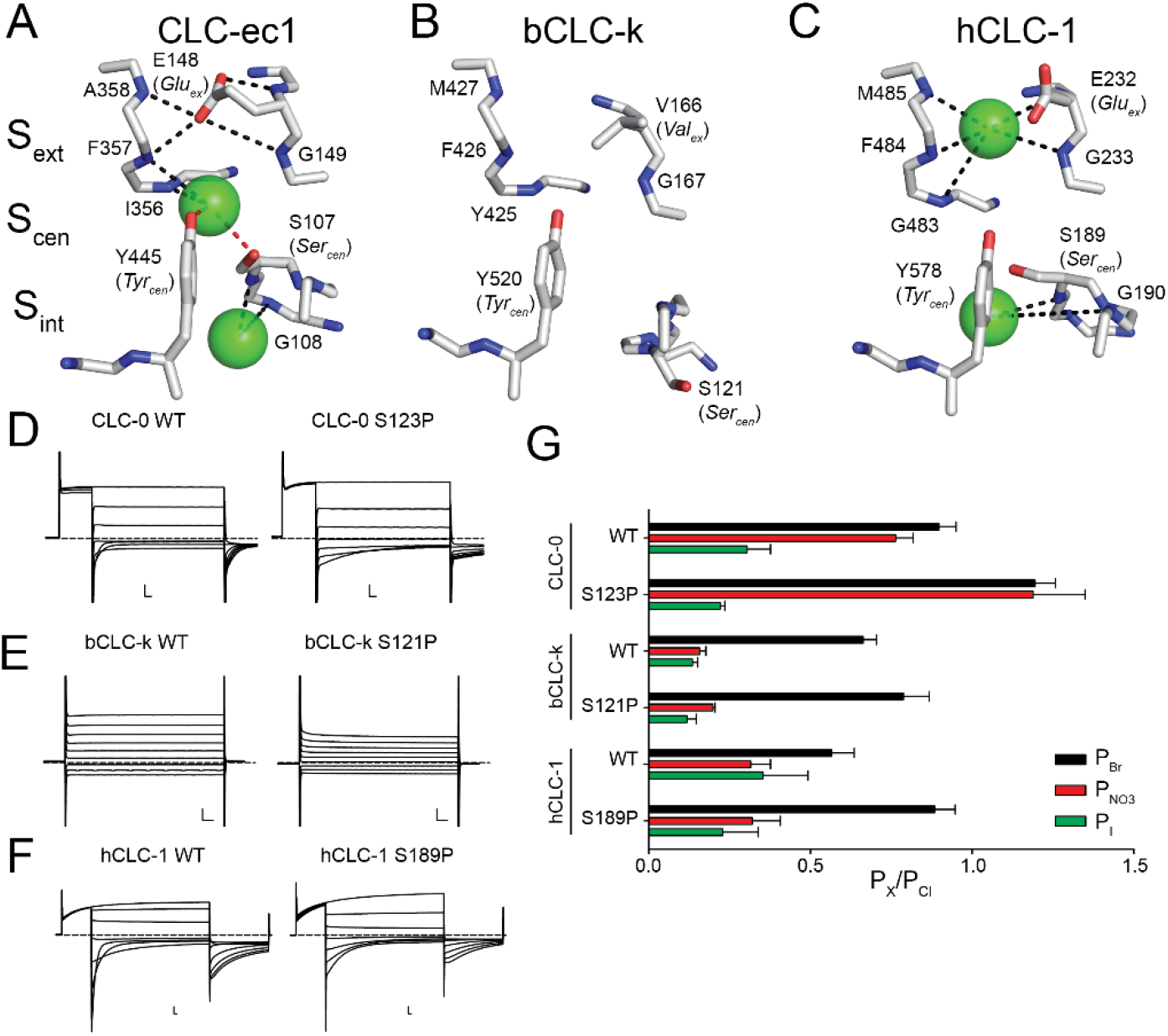
Structural architecture and ion coordination in the Cl^−^ pathway of CLC channels and transporters. (A-C) Close up view of the Cl^−^ permeation pathway in CLC-ec1 (PDB: 1OTS, A), bCLC-k (PBD: 5TQQ, B) and CLC-1 (PDB: 6COY, C). The position of the external (S_ext_), central (S_cen_) and internal (S_int_) binding sites is identified based on the crystal structure of CLC-ec1 ^31^. Bound Cl^−^ ions are shown as green spheres. No ions were resolved in the bCLC-k structure ^33^. Dashed lines indicate hydrogen bonds between the Cl^−^ ions and side chains (red) or backbone amides (black) ^31, 34^. D-F) Representative current traces of WT and proline mutants at Ser_cen_ in CLC-0 (D), bCLC-k (E) and hCLC-1 (F). Dashed lines indicate the 0 current level. Scale bars indicate 2 μA and 10 ms. G) The relative permeability ratios for Br^−^, NO_3_^−^ and I^−^ of CLC-0 (WT and S123P), bCLC-k (WT and S121P) and hCLC-1 (WT and S189P). Data are Mean ± S.E.M of n>7 repeats from N≥3 independent oocyte batches.

The current consensus mechanism for CLC selectivity is that the S_cen_ site is the primary regulator of anion discrimination and that the side chain of Ser_cen_ is the critical determinant of its specificity, as proline mutations at this site switch the selectivity from Cl^−^ to NO_3_^−^ and vice versa ^17, 19, 20, 21, 23, 50, 51, 52^. However, the recent finding that in bCLC-k channel Ser_cen_ points away from S_cen_^33^ (Fig. 1B) and that mutations at Ser_cen_ in the human CLC-Ka channel do not affect selectivity ^53^, recently led to the proposal that other pore-lining side chains are important for anion specificity. However, while side chains are not conserved, the functional preservation of the anion selectivity sequence points to a shared mechanism between hCLC-Ka and other CLC channels and transporters.

The extensive hydrogen bonding network of permeating anions with pore-lining backbone amides (Fig. 1A-C) led us to hypothesize that backbone amides might provide the conserved pattern of anion coordination in the CLCs, while side chain interactions contribute to fine-tuning of ion selectivity. Using a combination of atomic mutagenesis, electrophysiology, and molecular dynamics (MD) simulations we show that anion selectivity in CLC-0 and bCLC-k is determined by pore-lining backbone amides and their replacement with an ester oxygen destabilizes Cl^−^ binding with parallel effects on ion selectivity and permeation. Our results suggest that the role of S_cen_ and S_ext_ in ion selectivity depends on the side chain at the ‘gating glutamate’ position. When Glu_ex_ is a glutamic acid, selectivity is primarily determined at S_cen_, but when it is replaced with an uncharged (valine or alanine) residue, S_cen_ and S_ext_ play near-equivalent roles in ion selectivity. Our results shed new light onto the mechanism of anion permeation and selectivity in a CLC channel and show that backbone amides are critical in allowing these channels to specifically select Cl^−^ over other anions.

## Results

### C-D loop orientation does not determine the role of Ser_cen_ in anion selectivity

We tested whether the structural arrangement of the C-D loop (Fig. 1A-C) determines the role of Ser_cen_ in CLC selectivity by replacing this residue with a proline in CLC-0, CLC-1 and bCLC-k (Fig. 1D-G, Fig. 1 Supp. 2-3). Consistent with past results, the anion selectivity of CLC-0 is drastically altered by the S123P mutation, with the mutant becoming more permeable to Br^−^ and NO_3_^−^ than Cl^−^ (Fig. 1D, G, Fig. 1 Supp. 2-3) ^19, 21^. However, absent direct structural information on this channel it is difficult to interpret this effect. Thus, we introduced the corresponding mutation in the structurally known bCLC-k and hCLC-1 channels that differ in the orientation of Ser_cen_ (Fig. 1B-C, E-F, Fig. 1 Supp. 2-3) ^33, 34, 35^. Unlike S123P CLC-0, both constructs retain the selectivity sequences of their parent channels, Cl^−^>Br^−^>NO_3_^−^∼I^−^, with small alterations (Fig. 1G). Thus, the role of Ser_cen_ in CLC channel selectivity does not depend on the orientation of the C-D loop. This suggests other structural elements might play a more important role in the conserved selectivity sequence in CLCs.

### Backbone amides are key determinants of anion selectivity in bCLC-k and CLC-0

To test the role of backbone amides in anion selectivity we used the nonsense suppression method to site-specifically replace amino acids whose backbone amides may participate in ion coordination with their α-hydroxy acid equivalents ^54, 55^. This atomic manipulation “mutates” the peptide bond into an ester bond by substituting the backbone NH group with an oxygen atom (Fig. 2A), thus eliminating the backbone’s ability to function as an H-bond donor, without altering side-chain properties. The introduced ester is an otherwise modest change that shares similar bond lengths, angles, preference for a trans geometry, and comparably high energy barrier for rotation ^55, 56^. We chose the bCLC-k and CLC-0 channels as representatives CLC channels where Ser_cen_ does not (bCLC-k) or does (CLC-0) control anion selectivity. Incorporation of the α-hydroxy acids at the tested positions in bCLC-k and CLC-0 results in robust currents with measurable shifts in reversal potentials in different anions (Fig. 1 Supp. 2-3, Fig. 2 Supp. 1). The ratio of the currents measured in oocytes injected with tRNA conjugated to the UAA or with unconjugated tRNA is >9 at all positions (Fig. 3 Supp. 1A), suggesting that the contribution of currents due to non-specific incorporation and endogenous channels is ≤ 10%. The α-hydroxy acids for glycine or glutamate are not commercially available, and α-hydroxy substitutions in hCLC-1 did not yield sufficient currents for reversal potential determination.

**Figure 2.**
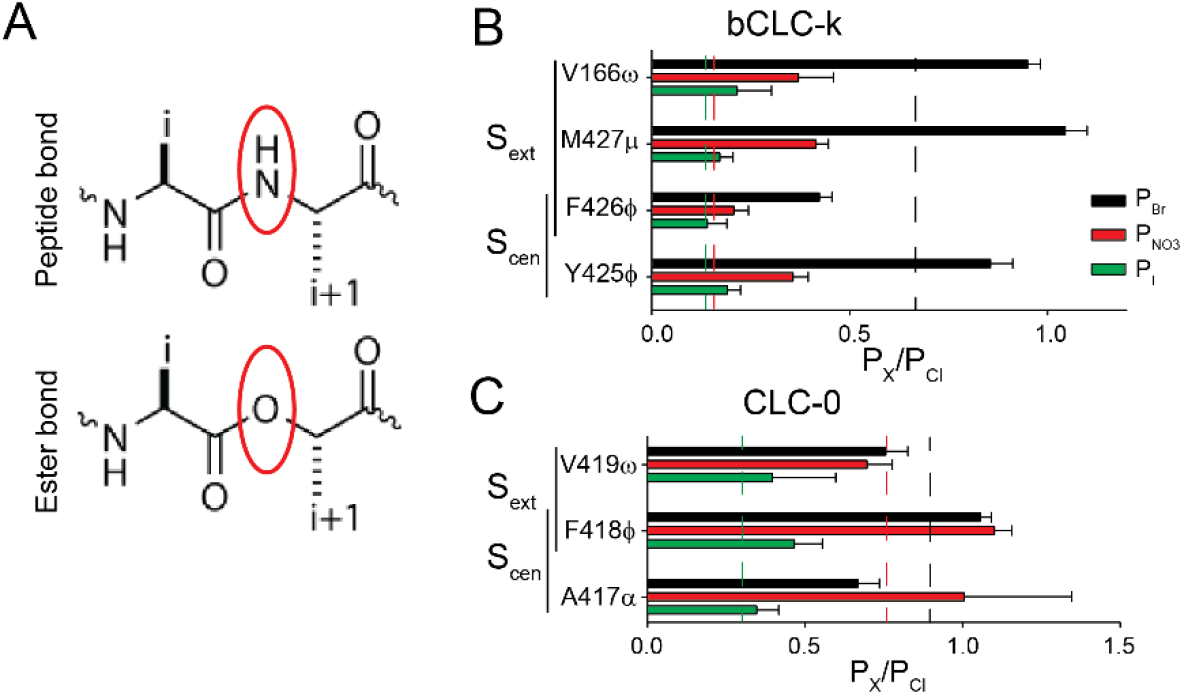
Role of backbone amides in anion selectivity of CLC-0 and bCLC-k. (A) Schematic representation of peptide (top panel) and ester bonds (bottom panel). (B-C) Effect of replacing backbone amides with ester oxygens at positions lining S_ext_ and S_cen_ in bCLC-k (B) and CLC-0 (C) on P_Br_ (black bars), P_NO_ (red bars) and P_I_ (green bars). Nomenclature of α-hydroxy acid substitutions is explained in Methods. All values are reported as mean ± S.E.M of n>7 repeats from N≥3 independent oocyte batches.

**Figure 3.**
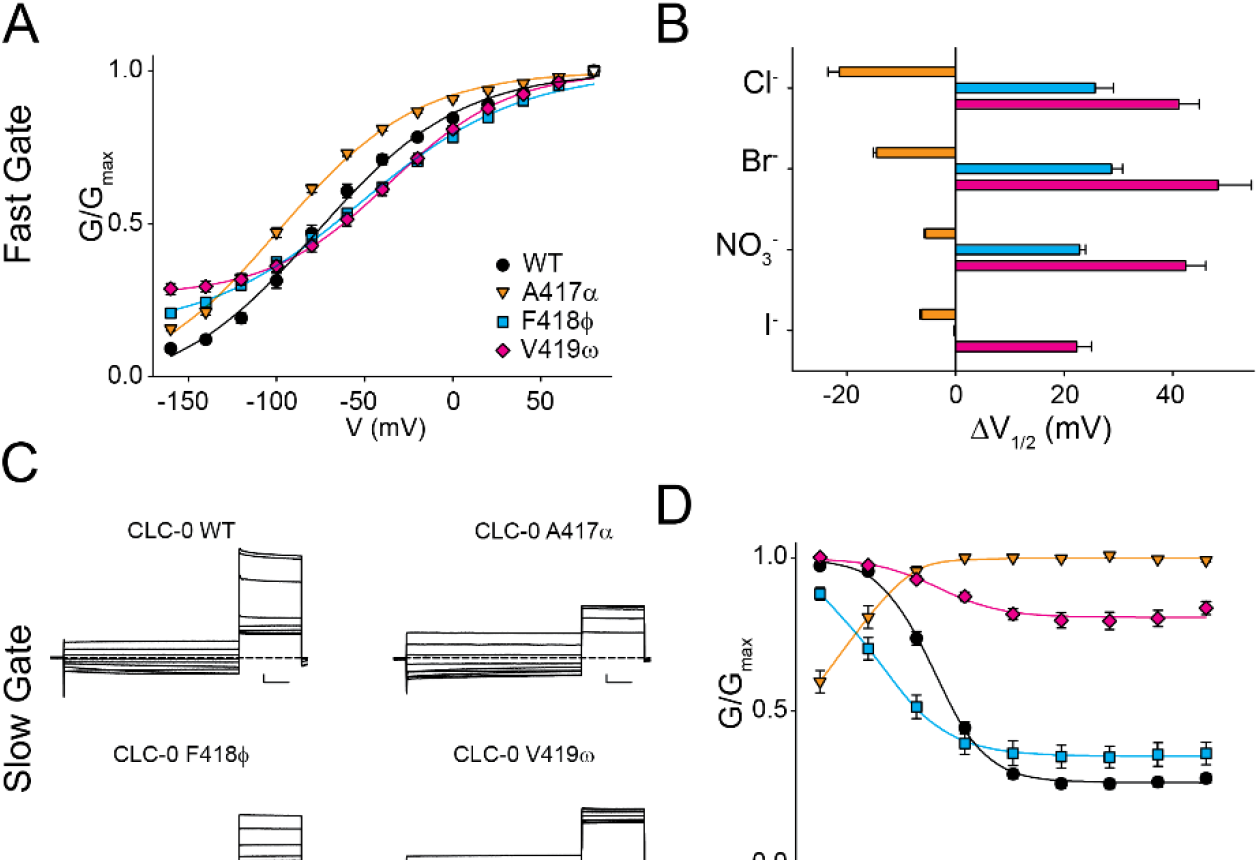
Pore-lining backbone amides affect fast and slow gating in CLC-0. A) Normalized G-V curves for fast gate of CLC-0 WT (black), A417α (orange), F418ϕ (cyan) and V419ω (pink) in Cl^−^. Solid lines are fits to Eq. 2. Values are mean ± S.E.M of n>7 repeats from N≥3 independent oocyte batches. B) ΔV_1/2_= (V_1/2_^mut^-V_1/2_^WT^) of the normalized fast gate G-V of WT and mutant CLC-0 in Cl^−^, Br^−^, NO_3_^−^ and I^−^. Colors as in A. Errors represent the propagation of the uncertainty of the V_1/2_ parameter evaluated from the fits of the data in Fig. 3A and Fig. 3 Supp 1. C) Representative slow gate current traces of WT and mutant CLC-0. Scale bars indicate 0.5 μA and 1 s. D) Normalized G-V curves for slow gate of CLC-0 WT (black), A417α (red), F418ϕ (green) and V419ω (yellow) in Cl^−^. Solid lines are fits to Eq. 2. Values are mean ± S.E.M of n>7 repeats from N≥3 independent oocyte batches.

In the bCLC-k channel, S_ext_ is lined by backbone amides of V166, M427 and F426, with F426 also lining S_cen_ together with Y425 (Fig. 1B). α-hydroxy substitutions at S_ext_, V166ω and M427μ, result in an altered selectivity order of Br^−^∼Cl^−^>NO_3_^−^>I^−^ while the others retain the WT order (Fig. 2B). The effects on P_Br_ and P_I_ are relatively small, <50% change relative to the WT values (Fig. 2 Supp. 2A) while effects on P_NO3_ are large at all positions, highlighted by a ∼250% P_NO3_ increase in M427μ (Fig. 2 Supp. 2A). Thus, backbone amides contribute to the overall selectivity of bCLC-k, and amides lining S_ext_ also control the inter-anionic selectivity sequence.

We used the same approach to investigate the selectivity of the CLC-0 channel and found that mutating backbones lining S_cen_ results in altered selectivity sequences, with F418ϕ not being able to discriminate between NO_3_^−^, Br^−^ and Cl^−^, and A417α showing an altered selectivity sequence of Cl^−^∼NO3^−^>Br^−^>I^−^ (Fig. 2C), while the S_ext_-lining V419ω substitution has a WT-like selectivity sequence (Fig. 2C). Thus, S_cen_ appears to primarily determine selectivity in CLC-0, consistent with previous results ^19, 21, 51^. Overall, the effects on P_Br_, P_NO3_ and P_I_ are relatively small, with <50% changes relative to the WT channel (Fig. 2 Supp. 2B), likely reflecting the weaker inter-anionic selectivity of CLC-0 compared to bCLC-k (Fig. 1G). In both channels, backbone substitutions have parallel effects on the permeability ratios and on the conductivity of the various ions, estimated from the ratio of the currents at +80 mV in the foreign anion to that of Cl^−^ (Fig. 2 Supp. 2C-H), indicating that interactions between backbone amides and the permeating ions determine binding and conduction.

### Pore-lining backbone amides play key roles in CLC-0 gating

Backbone mutations in the pore affect the G-V relationships of the single-pore gating process of CLC-0 in Cl^−^ (Fig. 3A, Fig. 3 Supp. 1), with the A417α substitution inducing a left shift in the G-V while the F418ϕ and V419ω replacements cause a right-shift in V_1/2_ (Fig. 3A-B). The direction of the V_1/2_ shifts is preserved for all mutants in Br^−^, NO_3_^−^ and I^−^, although the magnitudes vary (Fig. 3B). Correlation between the effects on selectivity and those on gating is poor, as the A417α mutation strongly alters selectivity while having comparatively modest effects on gating and, conversely, the V419ω replacement has a WT-like selectivity profile and the largest effects on the V_1/2_. Remarkably, these mutations have dramatic effects on the common-pore gating process (Fig. 3C), with A417α inverting its voltage dependence and V419ω resulting in a nearly constitutive phenotype (Fig. 3D). Thus, removal of a single hydrogen-bonding group in the CLC-0 pore affects the global rearrangements associated with slow gating ^57^, supporting the idea of a strong allosteric coupling between local and global rearrangements in this channel ^46, 47^.

### Glu_ex_ modulates the role of backbone-amides lining S_ext_ and S_cen_ in selectivity

The S_ext_ and S_cen_ sites play differential roles in determining selectivity of CLC-0 and bCLC-k channels (Fig. 2), despite the overall structural conservation of CLC pores (Fig. 1). One obvious difference between these channels is that the highly conserved Glu_ex_ of CLC-0 (E166) is replaced by an uncharged valine in bCLC-k (V166) (Fig. 1B, Fig. 1 Supp. 1E). Introducing Glu_ex_ in the bCLC-k channel (V166E) or eliminating it from CLC-0 (E166A), has minor effects on selectivity with the V166E bCLC-K channel maintaining a WT-like sequence of Cl^−^>Br^−^>NO_3_^−^∼I^−^ (Fig. 4A), while the E166A CLC-0 mutant has a slightly altered sequence of Cl^−^∼Br^−^≥NO_3_^−^>I^−^ (Fig. 4B).

**Figure 4.**
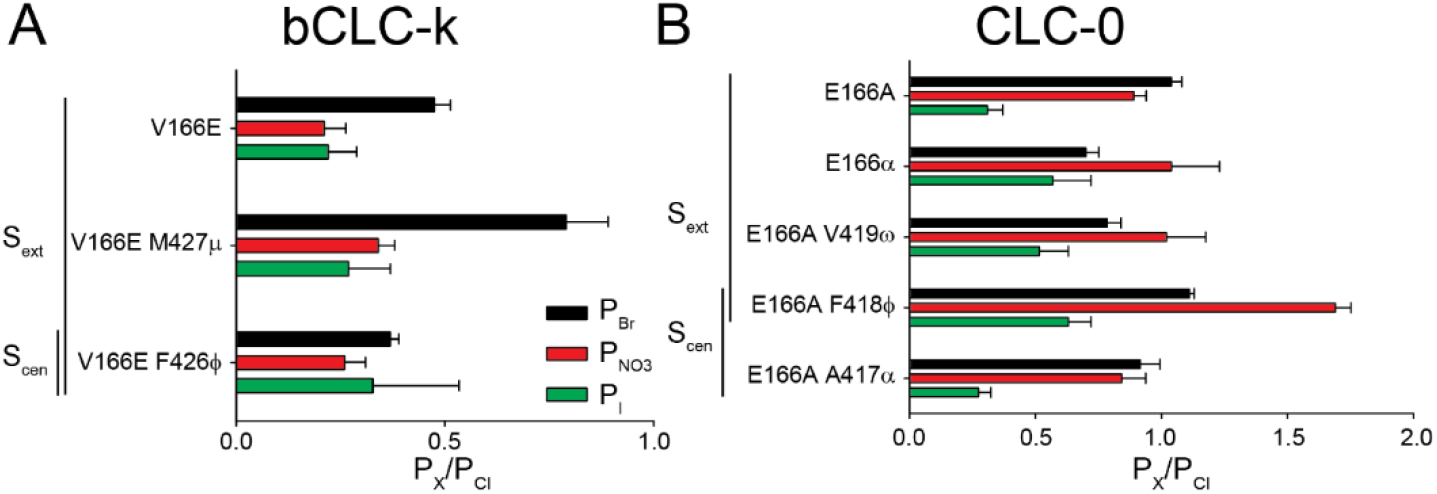
A glutamate side chain at the Glu_ex_ position modulates the role of S_cen_ and S_ext_ in anion selectivity. (A-B) Effect of replacing backbone amides with ester oxygens lining the S_ext_ and S_cen_ sites in V166E bCLC-k (A) and E166A CLC-0 (B) on P_Br_ (black bars), P_NO3_ (red bars) and P_I_ (green bars). All values are reported as mean ± S.E.M of N>7 repeats from at least 3 independent oocyte batches.

We then introduced the backbone ester substitutions on the background of these mutants to test how Glu_ex_ modulates the role of S_cen_ and S_ext_ in selectivity. Currents mediated by the V166E/Y425ϕ mutant were too small to obtain reliable results. The bCLC-k V166E/F426ϕ mutant, however, displays a marked preference for Cl^−^ and does not discriminate among other anions resulting in an altered selectivity sequence of Cl^−^>Br^−^∼I^−^∼NO_3_^−^ (Fig. 4A). In contrast, the V166E/M427μ mutant restores a WT-like selectivity sequence of Cl^−^>Br^−^>NO_3_^−^∼I^−^ (Fig. 4A), which was altered in the single M427μ mutant (Fig. 2B). In E166A CLC-0, the S_ext_-lining E166α and E166A/V419ω substitutions have an altered selectivity sequence of NO_3_^−^∼Cl^−^>Br^−^>I^−^, suggesting an increased role for S_ext_ in anion selectivity (Fig. 4B). Conversely, the S_cen_ lining E166A/A417α mutant restores a WT-like selectivity sequence (Fig. 4B) that was lost in the single A417α mutant (Fig. 2C). Most strikingly, the E166A/F418ϕ mutation converts CLC-0 into a NO_3_^−^-selective channel with a selectivity sequence of NO_3_^−^>>Br^−^>Cl^−^>I^−^ (Fig. 4B, Fig. 2 Supp. 2).

In summary, the energetic contribution of backbone amides lining S_ext_ increases in the absence of Glu_ex_, whereas, in S_cen_ only the contribution provided by F418 is influenced by Glu_ex_. Overall, backbone manipulation in the absence of Glu_ex_ has a larger relative effect on P_Br_, P_NO3_ and P_I_ than that in the presence of Glu_ex_ (Fig. 2 Supp. 2A-B). These results suggest that when Glu_ex_ is present S_cen_-lining backbone amides play a major role in anion selectivity, but when Glu_ex_ is replaced by a non-ionizable side chain selectivity is primarily determined by S_ext_-lining backbone amides. This difference likely reflects the ability of a glutamate side chain to compete with the permeant anions for occupancy of the S_cen_ site.

### Role of backbone amides in stabilizing anions in the bCLC-k pore

To investigate the contribution of the protein residues, particularly the backbone amides, to the binding of anions within the pore at a microscopic level, we simulated translocation of Cl^−^, Br^−^ or NO_3_^−^ through WT and M427*μμ* bCLC-k channels ^33^ and calculated the potential of mean force (PMF) profiles associated with these single-ion processes. For all anions, the PMF profiles show multiple local minima along the pore reporting on low-energy states of the ion where it establishes favorable interactions with the protein and local water molecules. Given the differences in sizes and H bonding patterns of the anions (Fig 5), small variations in the depth (∼1-2 kcal/mole) and the exact positions (∼1 Å) of these minima are observed.

**Figure 5.**
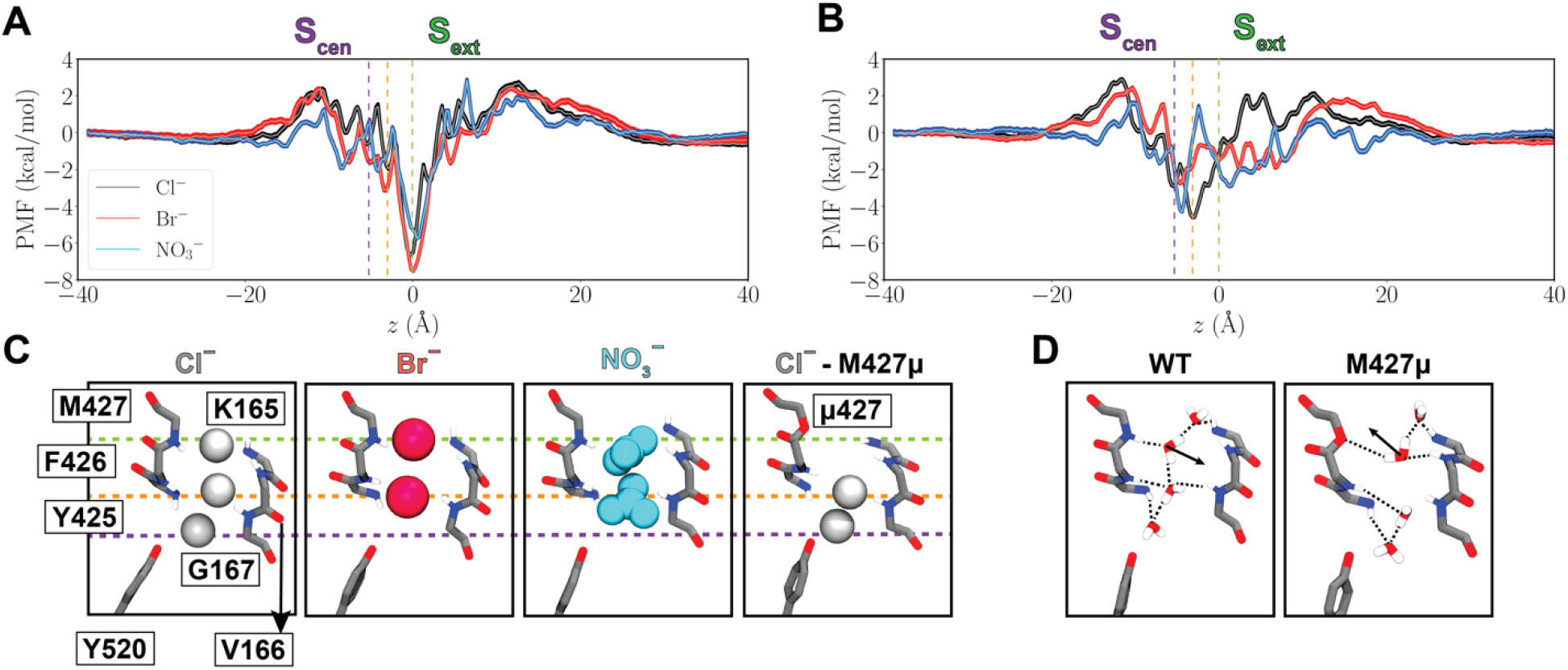
Anion-backbone interaction along the permeation pathway in WT and M427μ bCLC-k. (A) Single-occupancy PMF calculated along the ion permeation pathway for Cl^−^, Br^−^, and NO_3_^−^ in WT bCLC-k. The *z* positions for the ions are calculated relative to S_ext_ (*z* = 0). The positions of S_ext_, an intermediate binding region, and S_cen_ are indicated by the green, orange, and purple vertical dashed lines, respectively. PMFs are aligned by their energy in the bulk solution. (B) Single-occupancy PMF calculated along the ion permeation pathway for Cl^−^, Br^−^, and NO_3_^−^ in for M427μ bCLC-k. The uncertainty error in (B) and (C) is calculated based on Monte Carlo bootstrapping and shown in shaded colors. (C) The selectivity filter of bCLC-k along with the positions of all major binding sites (based on the PMFs, panels A and B) for different anions depicted in vdW for both WT and the M427μ mutant (Cl^−^ in white, Br^−^ in red, and NO_3_ ^−^ in blue). The three dashed lines correspond to those drawn in Panels A and B. (D) Comparison of hydration patterns in the selectivity filter of WT and M427μ bCLC-k, highlighting the shift in the orientations of water molecules.

The PMFs of WT channels show a global free energy minimum at S_ext_ for all three inspected anions (marked with a green dashed line in Fig. 5), highlighting this as the most stable anion-binding region along the pore. At this site the anions are mainly coordinated by the backbone amides of K165, V166, and M427. Among other minima, which are all significantly shallower than S_ext_, noticeable are the one at or around S_cen_ (marked with a purple dashed line in Fig. 5) in which the anions are coordinated by the backbone amide of Y425, and one in between S_ext_ and S_cen_ (marked with an orange dashed line in Fig. 5) where the anions are coordinated by the backbone amides of G167 and Y425. These results indicate that the backbone amide interactions play a key role in stabilizing the anions in all three minima (Fig. 5C). At all three sites, direct interactions between the anions and water molecules contributes to their stabilization. The M427μ replacement disrupts anion binding to S_ext_ (Fig. 5B) resulting in significant reduction of the well depth at this site or its complete disappearance as ions lose their favorable coordination with the amide hydrogen. Beyond local effects on S_ext_, this mutation also affects the pattern of hydration inside the constricted region, between S_ext_ and S_cen_, by altering the orientation of the water molecule coordinating M427 in the WT protein (Fig. 5D). Furthermore, our contact analysis on the permeant anions suggests that M427μ mutation affects the hydration pattern outside the constricted region (Fig. 5 Supp. 1A-B), which we believe is the reason for change in the PMF profiles observed in other regions of the pore. These results exemplify the critical contribution of backbone amide coordination for the anions within the pore. While this treatment does not represent the prevailing multi-ion permeation mechanism in these channels, it will probe the environment of the anions while in the pore and provides a reliable approximation for the energetics experienced by them.

## Discussion

Ion channels employ diverse molecular strategies to enable the rapid and selective passage of ions across biological membranes. While cation selectivity is relatively well understood, the molecular bases of anion selectivity remain poorly characterized. Indeed, many anion channels are more permeable to non-physiological ions, such as I^−^ or SCN^−^, than to the physiologically abundant Cl^−^ ^9, 10, 11, 12, 14^ and most small-molecule compounds designed to bind and transport anions share a similarly poor selectivity profile ^58, 59, 60^, highlighting our limited understanding of the fundamental mechanisms of anion selectivity. Unique among anion channels, the CLCs specifically select for Cl^−^ over other anions. Past work suggested that the CLC preference for Cl^−^ is determined by the specific interactions with a pore-lining serine side chain on the C-D loop, as substitutions at this position confer selectivity for other anions ^19, 20, 21, 23, 50, 52^. However, this mechanism was recently questioned as this loop adopts a different conformation in kidney-type CLC-k channels ^33, 53^ and its functional role is not conserved in the muscle-type CLC-1 channel (Fig. 1D). Thus, the determinants of the conserved Cl^−^ selectivity of the CLCs remain unknown.

Ions in the CLC permeation pathway form hydrogen bonds with backbone amides lining S_cen_ and S_ext_^33, 39^ whose position is well-conserved among CLC proteins (Fig. 1). We used atomic mutagenesis to site-specifically replace these putative hydrogen-bond donor backbone amides with an ester oxygen that cannot engage in hydrogen bonds with the permeating anions ^55, 56^. We found that targeted removal of individual pore-lining amides substantially degrades inter-anionic discrimination in both CLC-0 and bCLC-k, resulting in channels with weakened Cl^−^ preference (Fig. 2, 4). Indeed, elimination of a single hydrogen bond within the selectivity filter can increase P_NO3_ up to 250% and P_I_ up to 200% (Fig. 2 Supp. 4), while effects on P_Br_ are generally more modest, consistent with the idea that Br^−^ is a faithful substitute for Cl^−^ ^31, 38, 43, 48^. It is possible that the non-specific incorporation of amino acids and/or endogenous channels dampen the effects of backbone mutations on selectivity. However, currents recorded in oocytes injected with unconjugated tRNA are ≤ 10% of those from tRNA conjugated to the amino acids (Fig. 3 Supp. 1A), suggesting the error due to these contributions is relatively small and will not affect the trends of the effects we observe. It is possible that these errors could be higher for I^−^ currents, as this ion blocks CLCs but is more permeable through other Cl^−^ channels.

It is important to consider that ion permeation through the CLCs is a multi-ion process ^46^, which raises the possibility that ion selectivity could also be determined by multi-ion occupancy of the pore. However, several lines of evidence suggest this is not the case. First, the CLC channels and exchangers share the same selectivity sequence despite having different mechanisms of ion transport. Second, in CLC-ec1 −a CLC transporter that shares the same ion selectivity as the CLC-0 and bCLC-k channels studied here− the selectivity of ion binding and transport coincide ^18, 19^ and ion binding to the different sites is largely unaffected in single or multi-ion configurations ^19,38^. Finally, we showed that the effects of atomic-scale perturbations of individual sites can have profound effects on ion selectivity (Fig. 2, 4). Together, these observations suggest that the selectivity properties of the CLC Cl^−^ permeation pathway are primarily determined by the interactions of single ions with the pore and thus can be evaluated using single-ion PMF calculations (Fig. 5). Our free energy calculations highlight the involvement of backbone-ion coordination in the stabilization of the ions in the channel’s selectivity filter and in particular at the S_ext_ site, where we find that mutating the pore-lining backbone amide of M427μ directly and significantly destabilizes ion binding. Notably, our simulations show this manipulation also results in longer-range effects as it perturbs the hydration pattern of the pore (Fig. 5D). Finally, we found that substituting pore-lining backbone amides affects CLC-0 gating, with particularly marked effects on common gate activation. This supports the idea of tight allosteric coupling between ions permeating through the pore and the local and the global rearrangements that respectively underlie single- and common-pore gating processes in this channel ^46, 47^.

Our results show that in CLC-0 and CLC-k S_ext_ and S_cen_ play different roles, with selectivity in CLC-0 being primarily determined at S_cen_ while in bCLC-k both sites are important (Fig. 3, 4, 5). Although the structure of CLC-0 is unavailable, the orientation of backbone amides is well conserved in the pores of all available CLC channels and transporters (Fig. 1, Fig. 1 Supp 1), suggesting that it is unlikely that these functional differences arise from backbone structural rearrangements. Since in bCLC-k Glu_ex_ is replaced by an uncharged valine (Fig. 1B), we hypothesize these effects likely reflect the different interactions of a negatively charged or neutral side chain with the permeating ions as they compete for occupancy of the S_ext_ and S_cen_ sites. Indeed, removal of the Glu_ex_ side chain in CLC-0 increases the role of backbone amides lining S_ext_ in anion selectivity (Fig. 4B), and, conversely, introducing a glutamic acid in place of a valine in bCLC-k augments the role of S_cen_ in ion selectivity (Fig. 4A).

Our MD simulations suggest that in CLC-k the ions are most strongly bound at the S_ext_ site. In the backbone mutant M427μ, S_ext_ becomes unstable and the binding positions of ions and water near S_cen_ are shifted down with a clear change in their orientations, factors that likely result in the channel’s loss of selectivity for Cl^−^. The proposed mechanism, that ion selectivity is primarily determined via interactions with backbone elements, is reminiscent of the mechanism for selectivity in K^+^ channels ^1, 2^. The channel’s structure is optimized to provide an ideal coordination shell to the permeating ion via interaction with its backbone, where the choice of carbonyls or amides facilitates ions of different charge.

## Methods

### In vitro cRNA transcription

RNAs for all CLC-0 and bCLC-k wild-type and mutant constructs were transcribed from a pTLN vector using the mMessage mMachine SP6 Kit (Thermo Fisher Scientific, Grand Island, NY) ^46, 52, 61^. For final purification of cRNA the RNeasy Mini Kit (Quiagen, Hilden, Germany) was employed. RNA concentrations were determined by absorbance measurements at 260 nm and quality was confirmed on a 1% agarose gel.

### tRNA misacylation

For nonsense suppression of CLC-0 and bCLC-k TAG mutants in *Xenopus laevis* oocytes, THG73 and PylT tRNAs have been employed. THG73 was transcribed, folded and misacylated as previously described ^62^. PylT was synthetized by Integrated DNA Technologies, Inc. (Coralville, IA, USA), folded and misacylated as previously described ^63^. Ala-, Met-, Phe-, Val-, α-hydroxy Ala- (α), α-hydroxy Met- (μ), α-hydroxy Phe- (φ) and α-hydroxy Val-pdCpA (ω) substrates were synthesized according to published procedures ^63^.

### Nonsense suppression to replace amino acids with *α*-hydroxy acid

The nonsense suppression method to site-specifically replace amino acids with pore-lining backbone amides with their α-hydroxy acid equivalents ^55^. This atomic manipulation substitutes the backbone NH group with an oxygen atom, eliminating the ability of the backbone to function as H-bond donor without altering side chain properties (Fig. 4a), converting the peptide bond into an ester bond. These bonds have similar lengths, angles, preference for a trans geometry, and comparably high energy barrier for rotation ^55, 56^. Incorporation of the α-hydroxy acids at the positions tested in bCLC-k (V166, Y425, F426, M427) and CLC-0 (E166, A417, F418, V419) resulted in currents that were at least 5-fold higher than those recorded in oocytes injected with non-acetylated control tRNA (Fig. 3 Supp. 1A). We indicate mutations to α-hydroxy acids with their Greek letter counterpart: α for α-hydroxy alanine, ω for α-hydroxy valine, φ for α-hydroxy phenylalanine and μ α-hydroxy methionine. Incorporation of WT amino acids resulted in channels with WT-like properties (Fig. 3 Supp. 1B, C). Finally, insertion of φ at position F161 (F161φ) in CLC-0, a pore-lining residue located near Glu_ex_ (E166) but not involved in ion binding, resulted in WT-like selectivity (Fig. 3 Supp. 1D-E). These results indicate that effects on selectivity specifically reflect the incorporation of α-hydroxy acids at the targeted positions.

We were not able to test the role of the following residues: i) G164 (bCLC-k and CLC-0) because in our hands the α-hydroxy glycine acylated to the suppressor tRNA was extremely prone to hydrolysis impeding the incorporation; ii) K165 (bCLC-k) and R165 (CLC-0) because the corresponding α-hydroxy acids are not available; iii) E166 as α-hydroxy glutamic acid cannot be synthetized, and iv) we used α-hydroxy phenylalanine (φ) at position Y425 (bCLC-k) as the incorporation of α-hydroxy tyrosine was not successful. Phenylalanine was used as control in this case (Fig. 3 Supp. 1B). Currents associated with the V166E Y425φ mutant were too small to be analyzable. The nonsense suppression approach did not result in analyzable currents of CLC-1, CLC-5 or CLC-7.

### Protein expression in *Xenopus laevis* oocytes and two electrode voltage clamp (TEVC) recordings

*Xenopus laevis* oocytes were purchased from Ecocyte Bio Science (Austin, TX, USA) and Xenoocyte (Dexter, Michigan, USA) or kindly provided by Dr. Pablo Artigas (Texas Tech University, USA, protocol # 11024). For conventional CLC expression, the following injection and expression conditions have been used: for CLC-0, 0.1-5 ng cRNA were injected and currents were measured ∼6-24 h after injection; for CLC-1, ∼2 ng cRNA were injected and currents were measured ∼ 24 h after injection; ∼0.1 ng of each, CLC-K and Barttin cRNA, were coinjected and currents were measured the day after injection. For nonsense suppression of CLC-0 and bCLC-k constructs, cRNA and misacylated tRNA were coinjected (up to 25 ng of cRNA and up to 250 ng of tRNA per oocyte) and currents were recorded 6-24 h after injection.

TEVC was performed as described ^19, 64^. In brief, voltage-clamped chloride currents were recorded in ND96 solution (in mM: 96 NaCl, 2 KCl, 1.8 CaCl2, 1 MgCl2, 5 HEPES, pH 7.5) using an OC-725C voltage clamp amplifier (Warner Instruments, Hamden, CT). Ion substitution experiments were performed by replacing the 96 mM NaCl in the external solution with equimolar amounts of NaBr, NaNO_3_ or NaI. Data was acquired with Patchmaster (HEKA Elektronik, Lambrecht, Germany) at 5 kHz and filtered with Frequency Devices 8-pole Bessel filter at a corner frequency of 2 kHz. Analysis was performed using Ana (M. Pusch, Istituto di Biofisica, Genova), Sigmaplot (SPSS Inc.) and Prism (GraphPad, San Diego, CA, USA). For each substitution we recorded currents from oocytes injected with unconjugated tRNA (Fig. 3 Supp. 1A). This current, I(tRNA), reflects a combination of the contributions of CLC channels with non-specific incorporation of conventional amino acids and of the endogenous currents. In all cases, the ratio of the currents measured in oocytes injected with tRNA conjugated to the UAA, I(UAA), to I(tRNA) is >9 (Fig. 3 Supp. 1A). This suggests that the contribution of currents due to non-specific incorporation and endogenous channels is ≤10% and thus will not affect the trends of the observed effects. It is possible that the contribution to error could be higher for I^−^ currents, as this ion blocks CLCs but is more permeable through other Cl^−^ channels.

Oocytes were held at a resting potential of −30 mV. For CLC-0 two different recording protocols have been used to distinguish single-pore from common-pore gating. During the single-pore gating protocol the voltage was stepped to +80 mV for 50 ms and then a variable voltage from −160 mV to +80 mV increasing in 20 mV steps was applied for 200 ms, followed by a 50 ms pulse at −120 mV for tail current analysis. For CLC-0 common-pore gating, 7 s voltage steps from +20 mV to −140 mV have been applied in −20 mV increments followed by a 2.5 s +60 mV post pulse for tail current analysis. For bCLC-k the voltage was stepped to −30 mV for 20 ms and then a variable voltage from −80 mV to +80 mV increasing in 10 mV steps was applied for 150 ms, followed by a 20 ms pulse at −30 mV. For CLC-1 the voltage was stepped to +80 mV for 100 ms and then a variable voltage from −160 mV to +80 mV increasing in 20 mV steps was applied for 200 ms, followed by a 100 ms pulse at −100 mV for tail current analysis.

### Analysis of electrophysiological recordings

Permeability ratios were determined by measuring the change in reversal potential, ΔV_rev_, recorded upon substituting the external anion and using the Goldman-Hodgkin-Katz equation ^15^ as

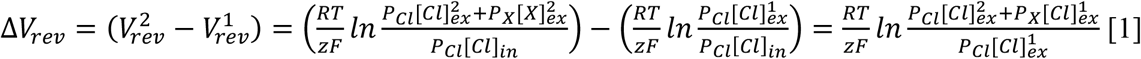

Where R, T, F and z have the usual meaning. The assumption that [Cl]_in_ did not change during successive perfusions was validated by bracketing recordings in Br^−^ and NO_3_^−^ with a return measurement in external Cl^−^ and ensuring that V_rev_ did not shift by more than 3 mV. Thus, the sequence of experiments was Cl^−^(1), Br^−^, Cl^−^(2), NO_3_^−^, Cl^−^ (3), I^−^ (Fig. 1 Supp. 3). In some cases, the order of Br^−^ and NO_3_^−^ was inverted, but no differences were detected. I^−^ was kept as the last ion tested due to its slow washout from oocytes. To simplify notation, throughout the text we indicate the relative permeability ratios of Br^−^, NO_3_^−^, and I^−^ as P_Br_, P_NO3_ and P_I_ with the understanding that these values represent the relative permeability ratios of these anions to that of Cl^−^, P_Br, NO3, I_/P_Cl_.

To estimate the voltage dependence of WT and mutant CLC-0, tail current analysis was performed, and data was fit to a Boltzmann function of the form:

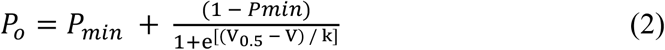

where *P*_*o*_ is the open probability as a function of voltage and is assumed to reach a value of unity at full activation. *P*_*min*_ is the residual open probability independent of voltage. V_0.5_ is the voltage at which 50% activation occurs, and k=RT/zF is the slope factor, R is the universal gas constant, T is temperature in K, F is the Faraday constant, and z is the gating charge.

### Statistical analysis

All values are presented as mean ± s.e.m. To determine statistical significance Student’s t-test (two-tailed distribution; two-sample equal variance) was performed. The threshold for significance was set to *P*=0.05.

### Simulation systems setup

For the bovine bCLC-k channel, the cryo-EM structures ^33^ (pdb:5TQQ) were used as the structural model for all the MD simulations and free energy calculations. Two unstructured loop regions missing in the cryo-EM structures (residue 258-276 and 454-456) were modeled using SuperLooper ^65^. The resulting models were embedded in lipid bilayers consisting of 80% POPC and 20% cholesterol and solvated with 0.15 M of NaCl and TIP3P water ^66^ using CHARMM-GUI MEMBRANE BUILDER ^67^. The dimension for the simulated systems was 150 × 150 × 130 Å^3^. The simulation system was energy-minimized for 10,000 steps, followed by two steps of 1-ns relaxation. The simulation system was then subjected to 1 ns of NPT initial equilibration with the standard protocol described in the CHARMM-GUI MEMBRANE BUILDER, which involves gradually releasing positional and dihedral restraints on the protein and lipid molecules. Thereafter, 10 ns of NPT equilibration with dihedral restraints (k = 100 kcal/mol/rad^2^) on the protein secondary structure were performed. After equilibration, the simulation system was equilibrated for another 250 ns without any restrains. This equilibrated system was used for all the free energy calculations.

### Simulation protocols

All simulations were carried out with NAMD 2.13 ^68, 69^, using CHARMM36m protein ^70^ and CHARMM36 lipid ^71^ parameters. The SHAKE algorithm ^72^ was employed to constrain bonds involving hydrogens to allow 2-fs timesteps for the integrator. A constant temperature of 310 K was maintained by Langevin thermostat ^73^ with a damping coefficient of 1 ps^−1^. Nosé-Hoover Langevin piston ^74^ with a period of 200 ps and a decay time of 50 ps was employed to maintain constant pressure at 1 atm. Periodic boundary conditions and a non-bonded cutoff of 12 Å (with a 10 Å switching distance and using vdW force switching) were used. Long-range electrostatics were calculated using the particle mesh Ewald method ^75^ with 1-Å grid spacing. Bonded interactions and short-range nonbonded interactions were calculated every timestep (2 fs). The pairs of atoms whose interactions were evaluated (neighborhood list) were updated every 20 fs. A cutoff (13.5 Å) slightly longer than the nonbonded cutoff was applied to search for interacting atom pairs.

### Free energy calculations

The free energy profiles, or the potential of mean force (PMF), of ion translocation through the permeation pore of bCLC-k were calculated using an enhanced sampling technique, umbrella sampling (US) ^76.^. In the US simulations, the reaction coordinate was chosen to be the *z* position (along the membrane normal) of the restrained ion relative to S_ext_ (set to *z* = 0), as the permeation pathway near the selectivity filter is roughly parallel to the membrane normal (aligned with the *z*-axis). To restrain the ion movement through to the selectivity filter, the *xy* coordinates of the ion were confined by a cylindrical half-harmonic wall (k = 10 kcal/mol/Å^2^) with a radius of 30 Å centering around the axis of the permeation pathway. For each ion the conduction pathway was divided into 80 umbrella windows with 1 Å interval and ranging from *z* = −40 Å to *z* = 40 Å, assuring the ion was in solution at ech end. In each window, the ion was harmonically restrained along the reaction coordinate (k = 5 kcal/mol/Å^2^), and initially equilibrated for 1 ns. Production sampling over each window was then done for 10 ns. The obtained distributions were then unbiased and combined using the weighted histogram analysis method (WHAM) ^77^ to obtain the PMF of the ion movement along the pore axis. The convergence of each PMF was examined by constructing the PMF after 5 to 10 ns of sampling with 1 ns interval. All the obtained PMFs remain unchanged after 9 ns of sampling, indicating a good degree of convergence (Fig. 5 Sup. 2). The final PMFs are constructed with 10 ns of sampling and uncertainty errors are calculated based on Monte Carlo bootstrapping.

## Data Availability

All constructs and electrophysiological traces are available on request.

## Statistics and Reproducibility

Functional experiments were repeated 7+ times from 3+ independent oocyte batches.

## Acknowledgements

The authors thank members of the Accardi lab for helpful discussions. The work was supported by National Institutes of Health (NIH) grants R01-GM128420 (to A.A.), R01-GM106569 and NINDS R24 NS104617 (to C.A.A.), R01-GM123455 and P41-GM104601 (to E.T.). Simulations in this study have been performed using allocations at National Science Foundation Supercomputing Centers (XSEDE grant number MCA06N060), and the Blue Waters Petascale Computing Facility of National Center for Supercomputing Applications (NCSA) at University of Illinois at Urbana-Champaign, which is supported by the National Science Foundation (awards OCI-0725070 and ACI-1238993) and the State of Illinois.

## Author contributions

L.L., K.L., S.D., C.A.A., E.T. and A.A. designed experiments; L.L., K.L., S.D., E.F. and J.G. performed experiments; L.L., K.L., S.D., E.T. and AA analyzed the data; A.A. prepared an initial draft and all authors edited the manuscript.

## References

1. Doyle DA, et al. The structure of the potassium channel: molecular basis of K^+^ conduction and selectivity. Science 280, 69–77 (1998).

2. Zhou Y, Morais-Cabral JH, Kaufman A, MacKinnon R. Chemistry of ion coordination and hydration revealed by a K^+^ channel-Fab complex at 2.0 Å resolution. Nature 414, 43–48 (2001).

3. Morais-Cabral JH, Zhou Y, MacKinnon R. Energetic optimization of ion conduction rate by the K+ selectivity filter. Nature 414, 37–42 (2001).

4. Noskov SY, Bernèche S, Roux B. Control of ion selectivity in potassium channels by electrostatic and dynamic properties of carbonyl ligands. Nature 431, 830–834. (2004).

5. Lockless SW, Zhou M, MacKinnon R. Structural and Thermodynamic Properties of Selective Ion Binding in a K+ Channel. PLoS Biology 5, e121 (2007).

6. Thompson A, Kim I, Panosian T, Iverson T, Allen T, Nimigean C. Mechanism of potassium-channel selectivity revealed by Na(+) and Li(+) binding sites within the KcsA pore. Nat Struct Mol Biol 16, 1317–1324 (2009).

7. Payandeh J, Scheuer T, Zheng N, Catterall WA. The crystal structure of a voltage-gated sodium channel. Nature 475, 353 (2011).

8. Tang L, et al. Structural basis for Ca2+ selectivity of a voltage-gated calcium channel. Nature 505, 56 (2013).

9. Bormann J, Hamill OP, Sakmann B. Mechanism of anion permeation through channels gated by glycine and gamma-aminobutyric acid in mouse cultured spinal neurones. The Journal of Physiology 385, 243–286 (1987).

10. Hwang T-C, Kirk KL. The CFTR ion channel: gating, regulation, and anion permeation. Cold Spring Harbor perspectives in medicine 3, a009498–a009498 (2013).

11. Ni Y-L, Kuan A-S, Chen T-Y. Activation and Inhibition of TMEM16A Calcium-Activated Chloride Channels. PLOS ONE 9, e86734 (2014).

12. Pifferi S. Permeation Mechanisms in the TMEM16B Calcium-Activated Chloride Channels. PLOS ONE 12, e0169572 (2017).

13. Vaisey G, Miller AN, Long SB. Distinct regions that control ion selectivity and calcium-dependent activation in the bestrophin ion channel. Proceedings of the National Academy of Sciences 113, E7399–E7408 (2016).

14. Qu Z, Hartzell C. Determinants of anion permeation in the second transmembrane domain of the mouse bestrophin-2 chloride channel. The Journal of general physiology 124, 371–382 (2004).

15. Hille B. Ion channels of excitable membranes, 3rd edn. Sinauer (2001).

16. Fahlke C, Dürr C, George AL, Jr. Mechanism of Ion Permeation in Skeletal Muscle Chloride Channels. Journal of General Physiology 110, 551–564 (1997).

17. Rychkov GY, Pusch M, Roberts ML, Jentsch TJ, Bretag AH. Permeation and block of the skeletal muscle chloride channel, ClC-1, by foreign anions. J Gen Physiol 111, 653–665 (1998).

18. Accardi A, Kolmakova-Partensky L, Williams C, Miller C. Ionic currents mediated by a prokaryotic homologue of CLC Cl^-^ channels. J Gen Physiol 123, 109–119. (2004).

19. Picollo A, Malvezzi M, Houtman JC, Accardi A. Basis of substrate binding and conservation of selectivity in the CLC family of channels and transporters. Nat Struct Mol Biol 16, 1294–1301 (2009).

20. Zifarelli G, Pusch M. Conversion of the 2 Cl(-)/1 H(+) antiporter ClC-5 in a NO(3)(-)/H(+) antiporter by a single point mutation. EMBO J 10, 1111–1116 (2009).

21. Bergsdorf EY, Zdebik AA, Jentsch TJ. Residues important for nitrate/proton coupling in plant and mammalian CLC transporters. J Biol Chem 284, 11184–11193 (2009).

22. De Angeli A, et al. The nitrate/proton antiporter AtCLCa mediates nitrate accumulation in plant vacuoles. Nature 442, 939–942. (2006).

23. Wege S, et al. The proline 160 in the selectivity filter of the Arabidopsis NO(3)(-)/H(+) exchanger AtCLCa is essential for nitrate accumulation in planta. Plant J 63, 861–869 (2010).

24. Brammer AE, Stockbridge RB, Miller C. F-/Cl-selectivity in CLCF-type F-/H+ antiporters. J Gen Physiol 144, 129–136 (2014).

25. Lim HH, Stockbridge RB, Miller C. Fluoride-dependent interruption of the transport cycle of a CLC Cl(-)/H(+) antiporter. Nat Chem Biol 9, 712–715 (2013).

26. Stockbridge RB, et al. Fluoride resistance and transport by riboswitch-controlled CLC antiporters. Proc Natl Acad Sci U S A 109, 15289–15294 (2012).

27. Last NB, et al. A CLC-type F-/H+ antiporter in ion-swapped conformations. Nature Structural & Molecular Biology 25, 601–606 (2018).

28. Jentsch TJ, Pusch M. CLC Chloride Channels and Transporters: Structure, Function, Physiology, and Disease. Physiological Reviews 98, 1493–1590 (2018).

29. Accardi A. Structure and gating of CLC channels and exchangers. The Journal of Physiology 593, 4129–4138 (2015).

30. Dutzler R. The structural basis of ClC chloride channel function. Trends Neurosci 27, 315–320 (2004).

31. Dutzler R, Campbell EB, MacKinnon R. Gating the selectivity filter in ClC chloride channels. Science 300, 108–112 (2003).

32. Feng L, Campbell EB, Hsiung Y, MacKinnon R. Structure of a Eukaryotic CLC Transporter Defines an Intermediate State in the Transport Cycle. Science 330, 635–641 (2010).

33. Park E, Campbell EB, MacKinnon R. Structure of a CLC chloride ion channel by cryo-electron microscopy. Nature 541, 500–505 (2017).

34. Park E, MacKinnon R. Structure of the CLC-1 chloride channel from Homo sapiens. eLife 7, e36629 (2018).

35. Wang K, et al. Structure of the human ClC-1 chloride channel. PLOS Biology 17, e3000218 (2019).

36. Schrecker M, Korobenko J, Hite RK. Cryo-EM structure of the lysosomal chloride-proton exchanger CLC-7 in complex with OSTM1. eLife 9, e59555 (2020).

37. Zhang S, et al. Molecular insights into the human CLC-7/Ostm1 transporter. Science Advances 6, eabb4747 (2020).

38. Lobet S, Dutzler R. Ion-binding properties of the ClC chloride selectivity filter. Embo J 25, 24–33 (2006).

39. Dutzler R, Campbell EB, Cadene M, Chait BT, MacKinnon R. X-ray structure of a ClC chloride channel at 3.0 Å reveals the molecular basis of anion selectivity. Nature 415, 287–294 (2002).

40. Accardi A, Miller C. Secondary active transport mediated by a prokaryotic homologue of ClC Cl^-^ channels. Nature 427, 803–807 (2004).

41. Miller C. ClC chloride channels viewed through a transporter lens. Nature 440, 484–489 (2006).

42. Feng L, Campbell EB, MacKinnon R. Molecular mechanism of proton transport in CLC Cl-/H+ exchange transporters. Proc Natl Acad Sci U S A 109, 11699–11704 (2012).

43. Nguitragool W, Miller C. Uncoupling of a CLC Cl^-^/H^+^ exchange transporter by polyatomic anions. J Mol Biol 362, 682–690 (2006).

44. Alekov AK, Fahlke C. Channel-like slippage modes in the human anion/proton exchanger ClC-4. J Gen Physiol 133, 485–496 (2009).

45. Orhan G, Fahlke C, Alekov AK. Anion- and proton-dependent gating of ClC-4 anion/proton transporter under uncoupling conditions. Biophys J 100, 1233–1241 (2011).

46. Pusch M, Ludewig U, Rehfeldt A, Jentsch TJ. Gating of the voltage-dependent chloride channel CIC-0 by the permeant anion. Nature 373, 527–531 (1995).

47. Chen TY, Miller C. Nonequilibrium gating and voltage dependence of the ClC-0 Cl^-^ channel. J Gen Physiol 108, 237–250 (1996).

48. Accardi A, Lobet S, Williams C, Miller C, Dutzler R. Synergism between halide binding and proton transport in a CLC-type exchanger. J Mol Biol 362, 691–699 (2006).

49. Walden M, Accardi A, Wu F, Xu C, Williams C, Miller C. Uncoupling and turnover in a Cl^-^/H^+^ exchange transporter. J Gen Physiol 129, 317–329 (2007).

50. Ludewig U, Pusch M, Jentsch TJ. Two physically distinct pores in the dimeric ClC-0 chloride channel. Nature 383, 340–343 (1996).

51. Ludewig U, Pusch M, Jentsch TJ. Independent gating of single pores in CLC-0 chloride channels. Biophys J 73, 789–797 (1997).

52. Leisle L, Ludwig CF, Wagner FA, Jentsch TJ, Stauber T. ClC-7 is a slowly voltage-gated 2Cl(-)/1H(+)-exchanger and requires Ostm1 for transport activity. EMBO J 30, 2140–2152 (2011).

53. Lagostena L, Zifarelli G, Picollo A. New Insights into the Mechanism of NO_3_^-^ Selectivity in the Human Kidney Chloride Channel ClC-Ka and the CLC Protein Family. Journal of the American Society of Nephrology 30, 293–302 (2019).

54. England PM, Zhang Y, Dougherty DA, Lester HA. Backbone mutations in transmembrane domains of a ligand-gated ion channel: implications for the mechanism of gating. Cell 96, 89–98 (1999).

55. Sereikaitë V, Jensen TMT, Bartling CRO, Jemth P, Pless SA, Strømgaard K. Probing Backbone Hydrogen Bonds in Proteins by Amide-to-Ester Mutations. Chembiochem : a European journal of chemical biology 19, 2136–2145 (2018).

56. Powers ET, Deechongkit S, Kelly JW. Backbone-Backbone H-Bonds Make Context-Dependent Contributions to Protein Folding Kinetics and Thermodynamics: Lessons from Amide-to-Ester Mutations. Advances in protein chemistry 72, 39–78 (2005).

57. Bykova EA, Zhang XD, Chen TY, Zheng J. Large movement in the C terminus of CLC-0 chloride channel during slow gating. Nat Struct Mol Biol 13, 1115–1119 (2006).

58. Wu X, et al. Nonprotonophoric Electrogenic Cl− Transport Mediated by Valinomycin-like Carriers. Chem 1, 127–146 (2016).

59. Hernando E, et al. Small molecule anionophores promote transmembrane anion permeation matching CFTR activity. Scientific reports 8, 2608 (2018).

60. Jentzsch AV, et al. Transmembrane anion transport mediated by halogen-bond donors. Nature Communications 3, 905 (2012).

61. Steinmeyer K, Schwappach B, Bens M, Vandewalle A, Jentsch TJ. Cloning and functional expression of rat CLC-5, a chloride channel related to kidney disease. J Biol Chem 270, 31172–31177 (1995).

62. Leisle L, et al. Cellular encoding of Cy dyes for single-molecule imaging. eLife 5, e19088 (2016).

63. Infield DT, Lueck JD, Galpin JD, Galles GD, Ahern CA. Orthogonality of Pyrrolysine tRNA in the Xenopus oocyte. Scientific reports 8, 5166–5166 (2018).

64. Leisle L, et al. Divergent Cl- and H+ pathways underlie transport coupling and gating in CLC exchangers and channels. eLife 9, e51224 (2020).

65. Hildebrand PW, et al. SuperLooper--a prediction server for the modeling of loops in globular and membrane proteins. Nucleic acids research 37, W571–574 (2009).

66. Jorgensen WL, Chandrasekhar J, Madura JD, Impey RW, Klein ML. Comparison of simple potential functions for simulating liquid water. J Chem Phys 79, 926–935 (1983).

67. Wu EL, et al. CHARMM-GUI Membrane Builder toward realistic biological membrane simulations. Journal of Computational Chemistry 35, 1997–2004 (2014).

68. Phillips JC, et al. Scalable molecular dynamics with NAMD. Journal of Computational Chemistry 26, 1781–1802 (2005).

69. Phillips JC, et al. Scalable molecular dynamics on CPU and GPU architectures with NAMD. The Journal of Chemical Physics 153, 044130 (2020).

70. Huang J, et al. CHARMM36m: an improved force field for folded and intrinsically disordered proteins. Nature Methods 14, 71–73 (2017).

71. Klauda JB, et al. Update of the CHARMM All-Atom Additive Force Field for Lipids: Validation on Six Lipid Types. Journal of Physical Chemistry B 114, 7830–7843 (2010).

72. Ryckaert J-P, Ciccotti G, Berendsen HJC. Numerical integration of the cartesian equations of motion of a system with constraints: molecular dynamics of n-alkanes. Journal of Computational Physics 23, 327–341 (1977).

73. Martyna GJ, Tobias DJ, Klein ML. Constant pressure molecular dynamics algorithms. The Journal of Chemical Physics 101, 4177–4189 (1994).

74. Feller SE, Zhang Y, Pastor RW, Brooks BR. Constant pressure molecular dynamics simulation: The Langevin piston method. The Journal of Chemical Physics 103, 4613–4621 (1995).

75. Darden T, York D, Pedersen L. Particle mesh Ewald: An N ·log(N) method for Ewald sums in large systems. The Journal of Chemical Physics 98, 10089–10092 (1993).

76. Kästner J. Umbrella sampling. WIREs Computational Molecular Science 1, 932–942 (2011).

77. Grossfield A. WHAM: the weighted histogram analysis method, version XXXX.

