## Supplementary Information for "Backbone amides are key determinants of Cl^−^ selectivity in CLC ion channels"

to

by

Lilia Leisle, Kin Lam, Sepehr Dehghani-Ghahnaviyeh, Eva Fortea, Jason Galpin, Christopher A. Ahern, Emad Tajkhorshid and Alessio Accardi

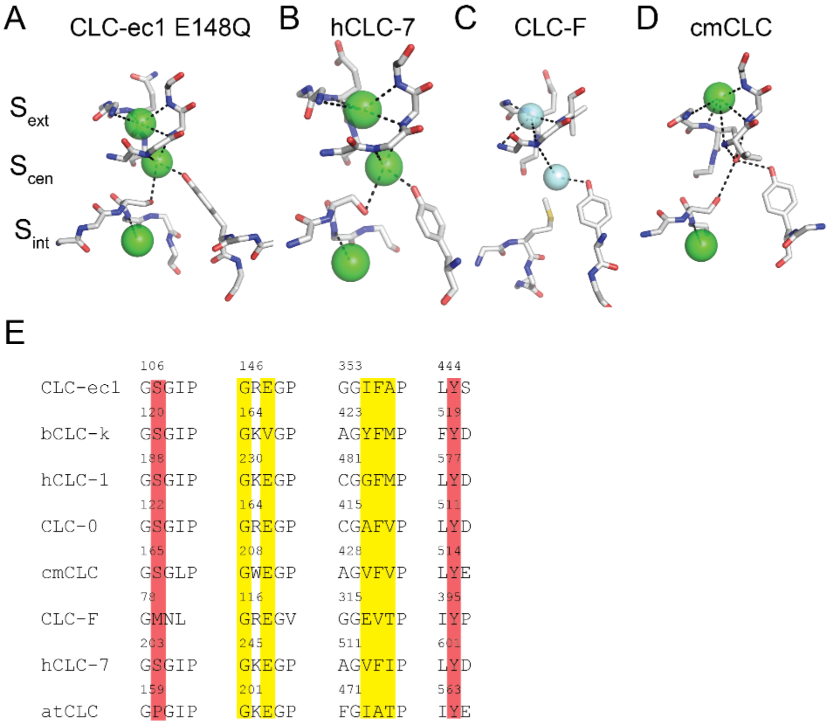


**Figure 1-Supplement 1. Structural and sequence conservation of the CLC ion pathway**. (A-D) Close up view of the $Cl^-$ permeation pathway in E148Q CLC-ec1(PDB: 1OTU, A), hCLC-7 (PDB: 7JM7, B), CLC-F (PDB: 6D0J, C) and cmCLC (PDB: 3ORG, D). Bound $Cl^-$ ($F^-$) ions are shown as green (cyan) spheres. Dashed black lines indicate hydrogen bonds between the anions and the protein. (E) Sequence alignment of the ion coordination motifs in CLC channels (bCLC-k, hCLC-1 and CLC-0) and transporters (CLC-ec1, hCLC-7, cmCLC, CLC-F, and atCLC-a). Yellow and red shading respectively indicate residues mediating anion coordination via backbone amides or side chains.

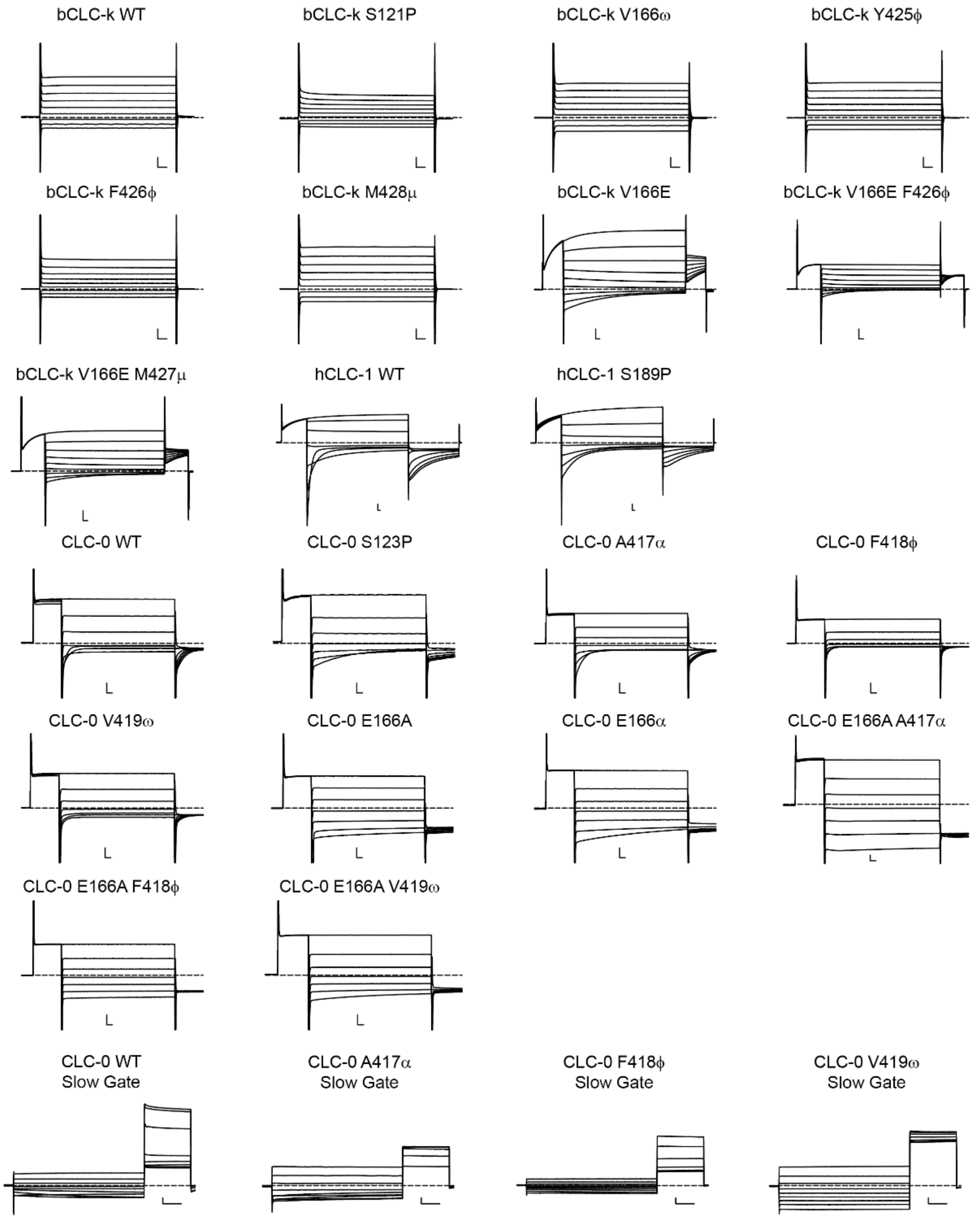


**Figure 1 Supplement 2. Representative currents of WT and all tested mutant constructs of bCLC-k, CLC-1 and CLC-0 channels.** Stimulation protocols are described in the methods section. Dashed lines indicate the 0 current level. In all cases scale bars indicate 2 μA and 10 ms apart from the bottom 4 panels where they correspond to 0.5 μA and 1 s.

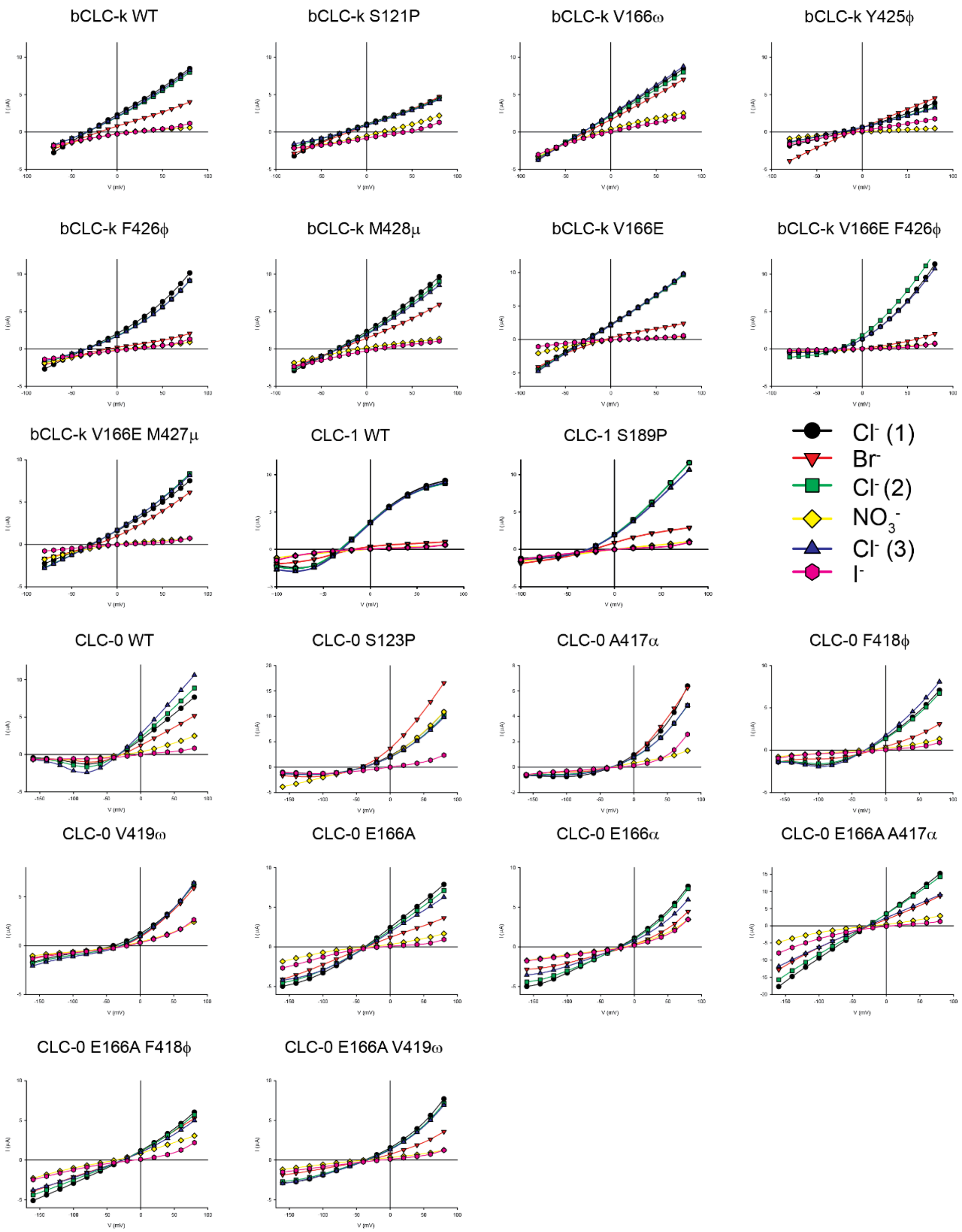


**Figure 1 Supplement 3. Representative I-V relationships for WT and all mutant constructs of bCLC-k, CLC-1 and CLC-0 channels.** Steady state I-V relationships measured during ion substitution experiments $Cl^-$(1) (black circles), $Br^-$ (red triangles), $Cl^-$(2) (green squares), $NO_3^-$ (yellow diamond), $Cl^-$(3) (blue inverted triangle), $I^-$ (pink hexagon). Experiments shown are for a single cell and un-normalized.

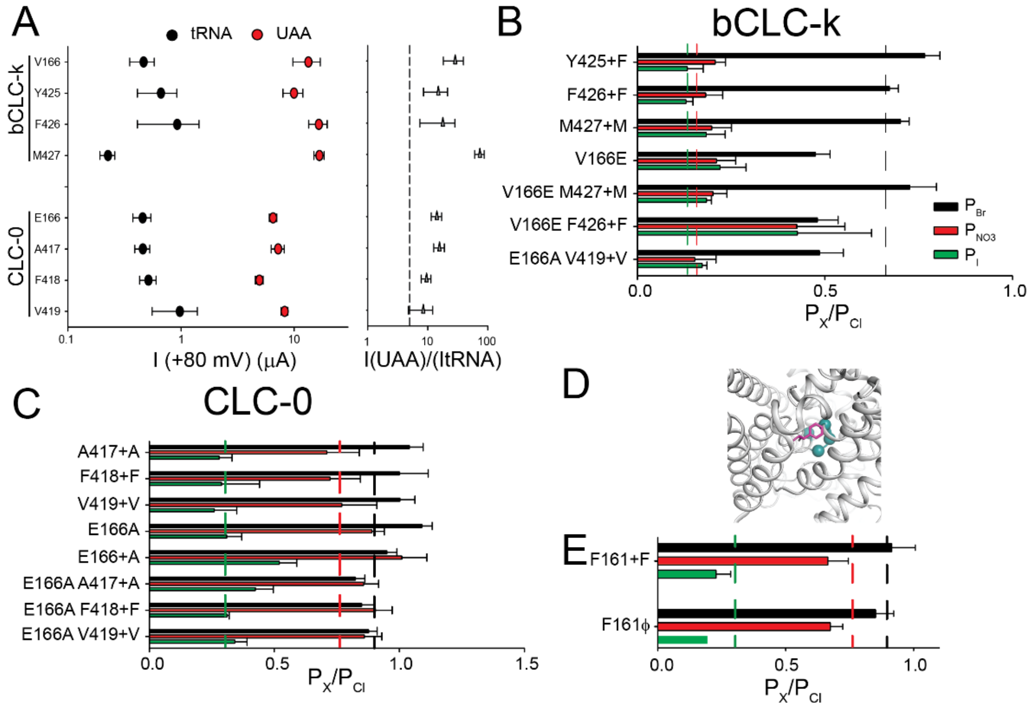


**Figure 2-Supplement 1 Application of the nonsense suppression method to CLC $Cl^-$ channels** (A) (right panel) Site-specific incorporation of non-canonical amino acids into the bCLC-k and CLC-0 channels is efficient and yields robust currents. Mean current I at +80 mV in *Xenopus laevis* oocytes for bCLC-k V166X, Y425X, F426X and M427X and CLC-0 E166X, A417X, F418X, V419X co-injected with empty tRNA (black) or misacylated tRNA loaded with the corresponding α-hydroxy acid (red). (left panel) The ratio I(UAA)/I(tRNA) was calculated from the mean values of I(UAA) and I(tRNA) at +80 mV. Errors were propagated. A threshold of I(UAA)/I(tRNA) >5 is imposed for specific incorporation efficiency (dashed line). (B-C) Incorporation of conventional aminoacids in bCLC-k (B) and CLC-0 (C) channels using the nonsense suppression method results in WT-like ion selectivity profiles. (D) Close up view of the outer vestibule of the bCLC-k pore highlighting the position of F161 (pink stick), and the Cα atoms of V166, Y425, F426 and M427 (cyan spheres). (E) Incorporation of Phe or φ at position F161 in CLC-0 results in WT-like ion selectivity. All values are reported as mean ± S.E.M of N>7 repeats from at least 3 independent oocyte batches.

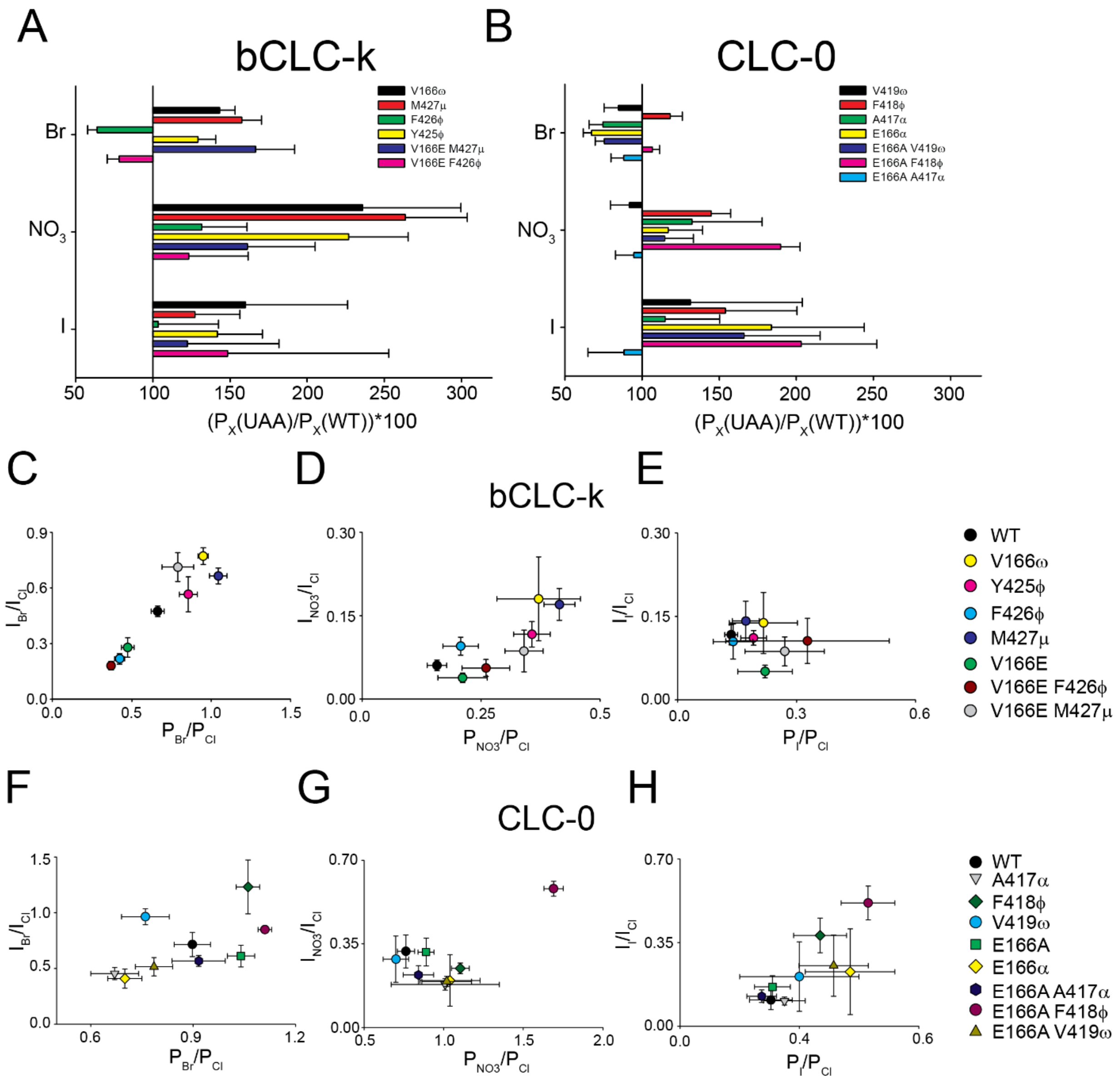


**Figure 2 Supplement 2. Effects of backbone amide substitutions on ion permeability and ion conductivity.** A-B) Normalized effects on $P_{Br}$, $P_{NO3}$ and $P_{I}$ elicited by backbone mutations in bCLC-k (A) and CLC-0 (B). To compare effects due to backbone substitutions, values of single mutants were normalized to those of the WT parent channel, and those in the $Glu_{ex}$ mutant background were normalized to those of the parent single mutant, V166E for bCLC-k and E166A for CLC-0. C-H) Relationship between the effects of backbone mutations in bCLC-k (C-E) and CLC-0 (F-H) on permeability and conduction of $Br^-$ (C, F), $NO_3^-$ (D, G) and $I^-$ (E, H).

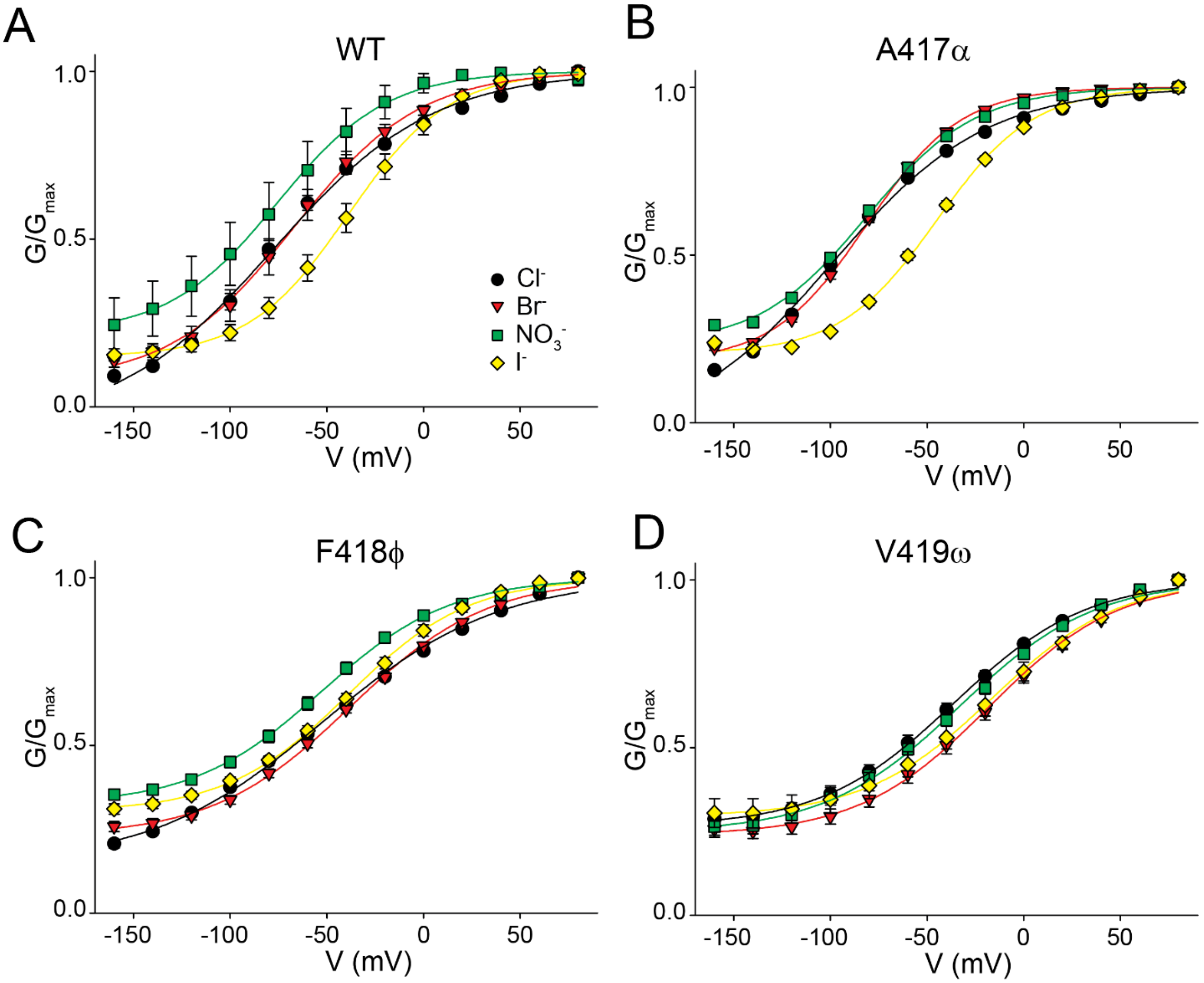


**Figure 3 Supplement 1. Backbone mutations affect the anion-dependent modulation of CLC-0 fast gating.** A-D) Average fast gate G-V relationships for WT (A), A417α (B), F418φ (C) and V419ω (D) measured in $Cl^-$ (black circles), $Br^-$ (red triangles), $NO_3^-$ (green squares) and $I^-$ (yellow diamond).

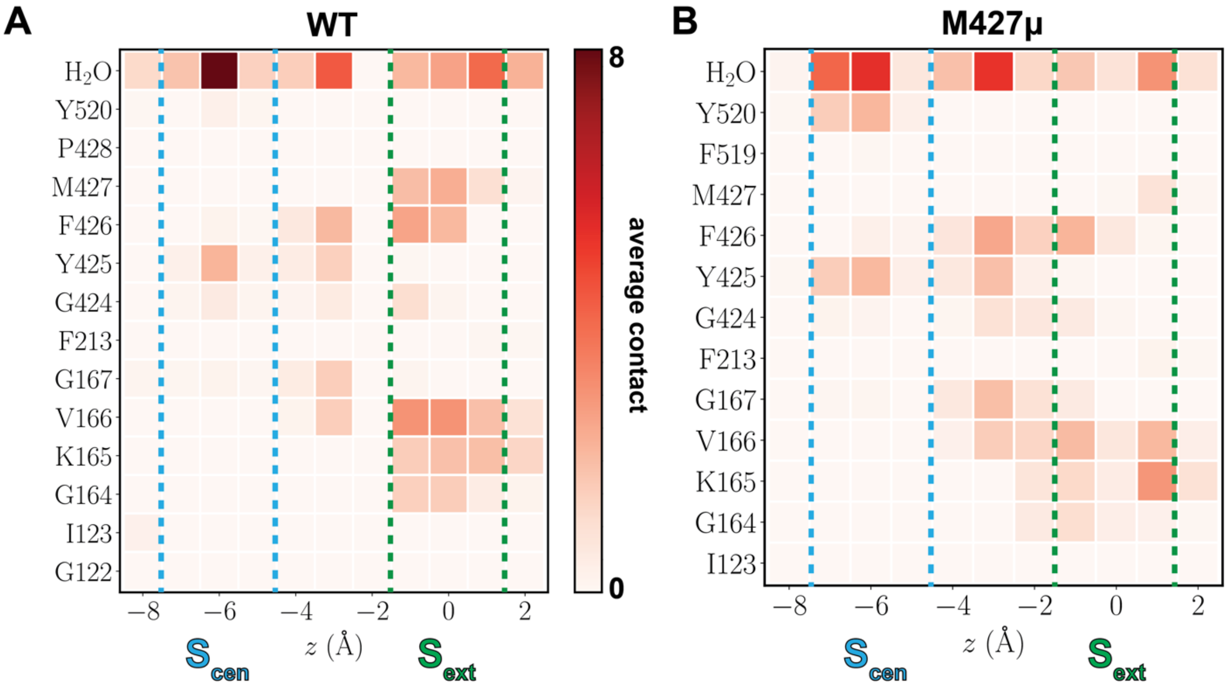


**Figure 5-Supplement 1. Average number of contacts of a $Cl^-$ ion at $S_{cen}$ and $S_{ext}$ sites for WT and M427μ bCLC-k.** (A) and (B) $Cl^-$ contacts with hydrogen atoms of the protein residues and water molecules for WT and M427μ bCLC-k, respectively. The contacts are calculated when the ion is located between z = -8 Å to z = 2 Å ($S_{ext}$ is at z = 0). A hydrogen atom is considered to make a contact if it is located within 3 Å of the ion. Boundaries of $S_{cen}$ and $S_{ext}$ sites are shown with dashed cyan and green lines, respectively. The colormap corresponds to the number of contacts throughout the trajectory with minimum and maximum number of contacts shown with light and dark red, respectively. The water contacts are changed due to the mutation at $S_{ext}$, suggesting a different hydration pattern upon the mutation.

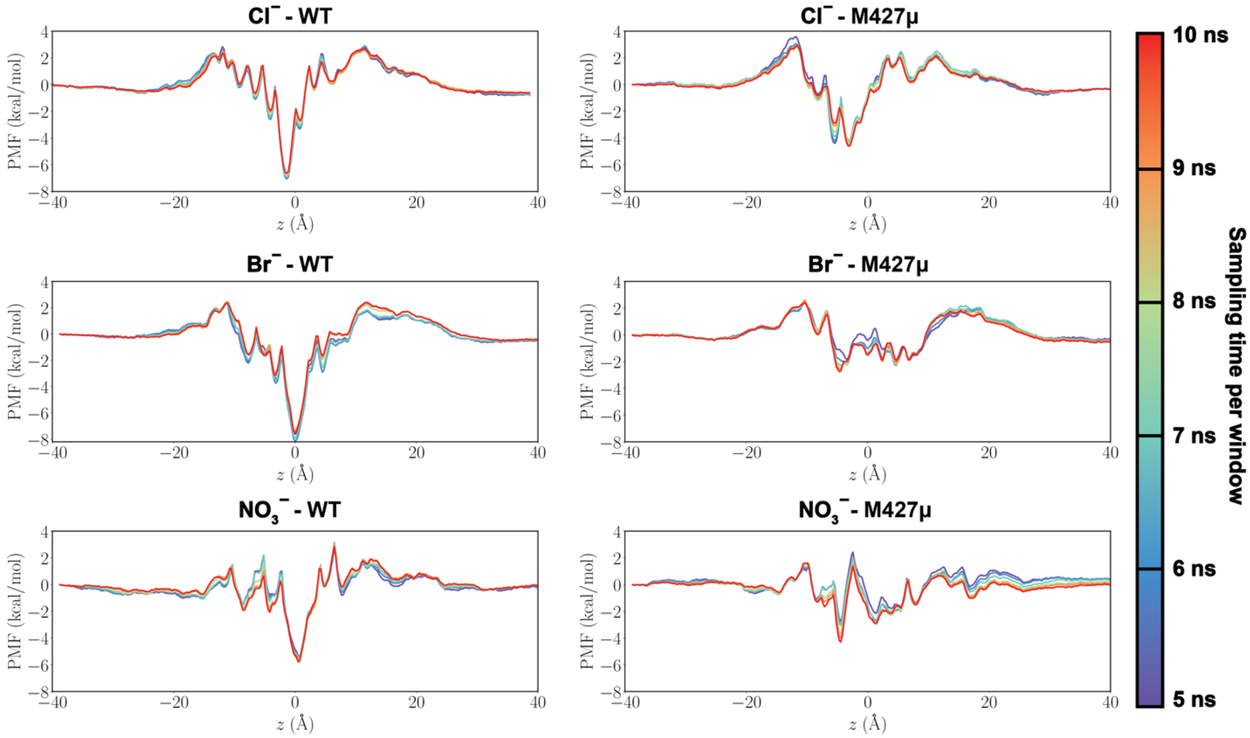


**Figure 5-Supplement 2. Examining the convergence of each PMF curve.** To examine the convergence of the PMFs, each curve is constructed after 5 ns (blue) to 10 ns (red) of sampling with 1 ns interval, for both WT and M427μ systems. The PMFs remain unchanged after 9 ns of sampling.
